# OMNI: Optimized Multi-view Network Integration with Heterogeneous Graph Attention for Biomedical Interaction Prediction

**DOI:** 10.64898/2025.12.25.696543

**Authors:** Gori Sankar Borah, Sukriti Tiwari, Selvaraman Nagamani

## Abstract

**Motivation:** Accurate prediction of biomedical relationships, such as chemical–gene interactions, is fundamental to understanding disease mechanisms and advancing drug discovery. With the rapid growth of heterogeneous biological data, modeling large-scale, multi-entity networks has become increasingly challenging. Traditional approaches, including homogeneous GNNs (e.g., GCN, GAT) and meta-path-based random walks, struggle to efficiently capture high-order, diverse neighborhood information in complex biomedical graphs. To address these limitations, we propose a novel multi-view heterogeneous graph attention network (GAT)-based architecture that effectively aggregates rich, heterogeneous interactions across multiple biomedical entity types. The proposed encoder captures comprehensive structural and semantic information while remaining computationally efficient. Through optimized aggregation strategies and multi-processing, the model generates high-quality node embeddings with significantly reduced training time. For relation prediction, multiple decoder architectures were evaluated, with a multilayer perceptron (MLP) identified as the most effective for accurate multi-type relation classification. The resulting network comprises 124,604 unique nodes and 48,482,286 interactions.

**Results:** Experimental results show that the proposed model consistently outperforms state-of-the-art methods, including CGINet, Node2Vec, and the GCN-based BioNet, achieving an AUROC of 0.91 for chemical–gene interaction prediction. The model further explores its ability to identify top-ranking chemical-gene interactions in cancer and to predict gene-phytochemical relationships. Overall, this work introduces a scalable and powerful framework for biomedical relation prediction, with strong potential applications in drug screening and disease mechanism discovery.

**Key Points:**

- We constructed a large-scale heterogeneous biological interaction network by integrating curated datasets across multiple entity types, including chemicals, genes, pathways, and diseases.
- We propose a novel graph neural network framework, Optimized Multi-View Network Integration (OMNI), based on an encoder–decoder architecture, which employs a multi-view heterogeneous Graph Attention Network (GAT) to learn entity embeddings from subgraphs and a multilayer perceptron (MLP) decoder to predict chemical–gene interactions (CGIs).
- We integrated a *PyTorch Lightning* based parallel training strategy to scale up the learning process, significantly enhancing the model’s ability to efficiently handle large-scale heterogeneous data.
- We demonstrated the applicability of OMI by evaluating cancer-related chemical–gene interactions and vitamin D receptor (VDR)–phytochemical interactions, including the prediction of interaction types.

## Introduction

The traditional drug discovery process is a time-consuming and resource-intensive approach, and it takes at least a decade to bring a drug to the market [1]. Recent advancements in computational power and novel algorithms have greatly revolutionised drug discovery methods, as well as drug repurposing strategies. These strategies substantially reduce the cost and time required for therapeutic discovery compared to traditional methodologies [2]. Most drugs work on the principle that a small molecule targets a single/multiple protein targets to achieve therapeutic effects [3]. Analysing and uncovering unknown CGIs provides a unique opportunity in the drug discovery field [5–7]. High-throughput screening, gene expression profiling, proteomics, metabolomics, and computational methods (i.e., structure- analogue-based approaches, network analysis) are some of the popular approaches available to study chemical-gene interactions (CGIs) [4]. Despite their utility, the heterogeneity, rapid growth, and high dimensionality of biological data pose substantial challenges for comprehensive CGI analysis. Importantly, many existing approaches focus predominantly on direct CGIs, often overlooking the broader biological context in which these interactions operate.

The interdependent and intricate relationships among different biological entities underscore the need for a more integrative analytical framework. Thus, incorporating additional biological entities, such as biological pathways and diseases, enables a systems-level understanding of CGIs. Several studies were conducted by collating a large-scale heterogeneous biological interaction network extracted from the literature. Different methodologies, such as network pharmacology [8,9], were applied to analyse this huge network. However, the rapid expansion of available data and the limited analytical methods face challenges in processing, integrating and extracting meaningful insights. In this context, data-driven approaches, including ML and systems-level modeling, have emerged as powerful tools for analyzing and interpreting massive biological networks.

Link prediction between biological entities can be effectively handled by various DL methods, especially graph-based models. The relation prediction model can extract and decode new relations by characterising known interrelations among biological entities. Li et. al. applied a graph neural network (GNN) for drug repurposing strategies by decoding the relationships among diseases, targets and drugs (DTD-GNN) [10]. Different graph network architectures, such as GNN [11], graph convolutional neural network (GCN) [12], and graph attention network (GAT) [13], were applied to decode drug-disease association prediction. BiopathNet [14] is a novel graph network architecture for link prediction based on path-based reasoning.

These traditional and advanced architectures are novel and unique in their ability to predict CGIs. However, they have several limitations, i.e., 1. Some of the methods have poor prediction accuracy due to their inability to merge and embed information from adjacent nodes. 2. The reliance on limited scale or incomplete datasets limits the ability to capture the full spectrum of biological relationships, thereby restricting the relationship prediction performance.

Recent advancements in biological databases and novel algorithms offer an opportunity to develop more reliable graph-based models with enriched data, facilitating the training of large-scale networks. For instance, BioNet [14] is an algorithm with a novel encoder-decoder architecture and an applied GCN for interaction prediction from a large-scale heterogeneous biological network. However, these types of models have different limitations, including 1. Methods that emphasize direct interactions in node representation learning often weaken the contribution of distant nodes, leading to information loss across the network 2. Many heterogeneous graph neural networks (HGNNs) employ random walk strategies to capture additional contextual information. However, they often underperform compared to methods that incorporate information from all neighbouring nodes in link prediction tasks 3. BioNet applies parallel computation to enhance learning efficiency, but its complexity poses challenges for biological scientists to adopt and reproduce it for their research. In this study, we propose a scalable modified multi-view heterogeneous GAT to predict the CGIs. The major contributions of this work are as follows,

1. A large-scale heterogeneous biological interaction network was constructed by integrating curated datasets from different entities, chemicals, genes, pathways, and diseases.
2. A novel GNN model named OMNI (Optimized Multi-view Network Integration) is proposed based on an encoder-decoder architecture [15] that utilizes a multi-view heterogeneous graph attention network to learn entity embeddings from subgraphs and different decoders (i.e. Multi Layer Perceptron (MLP) [16], MultX [17], TransE [18], Reverse [19]) were benchmarked to find a suitable decomposition decoder to predict CGIs. The existing architecture is modified and simplified to effectively handle massive data and create embeddings in less time without losing data information.
3. We integrate ‘*PyTorch Lightning*’ [20], a DL framework, to scale up the performance of OMNI and improve the model’s ability to handle large-scale data.
4. The robustness of the developed model has been demonstrated by evaluating its potential in the prediction of cancer-related CGIs and phytochemicals – Vitamin D

Receptor (VDR) interaction to prioritizes chemicals with higher potential for effective therapeutics.

## Materials and methods

### OMNI model architecture

A graph (G) can be denoted as (V, E). Here, V is the given set of nodes V={𝑣_𝑖_}, E is the set of edges *E=*{(𝑣_𝑖_, *r*, 𝑣_𝑗_)}, the r represents the type of edge. The objective of our model is to estimate the probability of a target edge 𝑒_𝑖𝑗_ = (𝑣_𝑖_, *r*, 𝑣_𝑗_). The OMNI adopts a novel encoder-decoder architecture to calculate this probability. The encoder leverages an optimized multi-view heterogeneous GAT to learn node representations. Different decoders (i.e. MLP, DistMult, ComplEX, RESCAL) were benchmarked to identify the potential decoder that can effectively predict the likelihood of candidate edges. **Figure 1** represents the panorama of OMNI, where the input is a graph structure and nodes are represented as an adjacency matrix.

**Figure 1.**
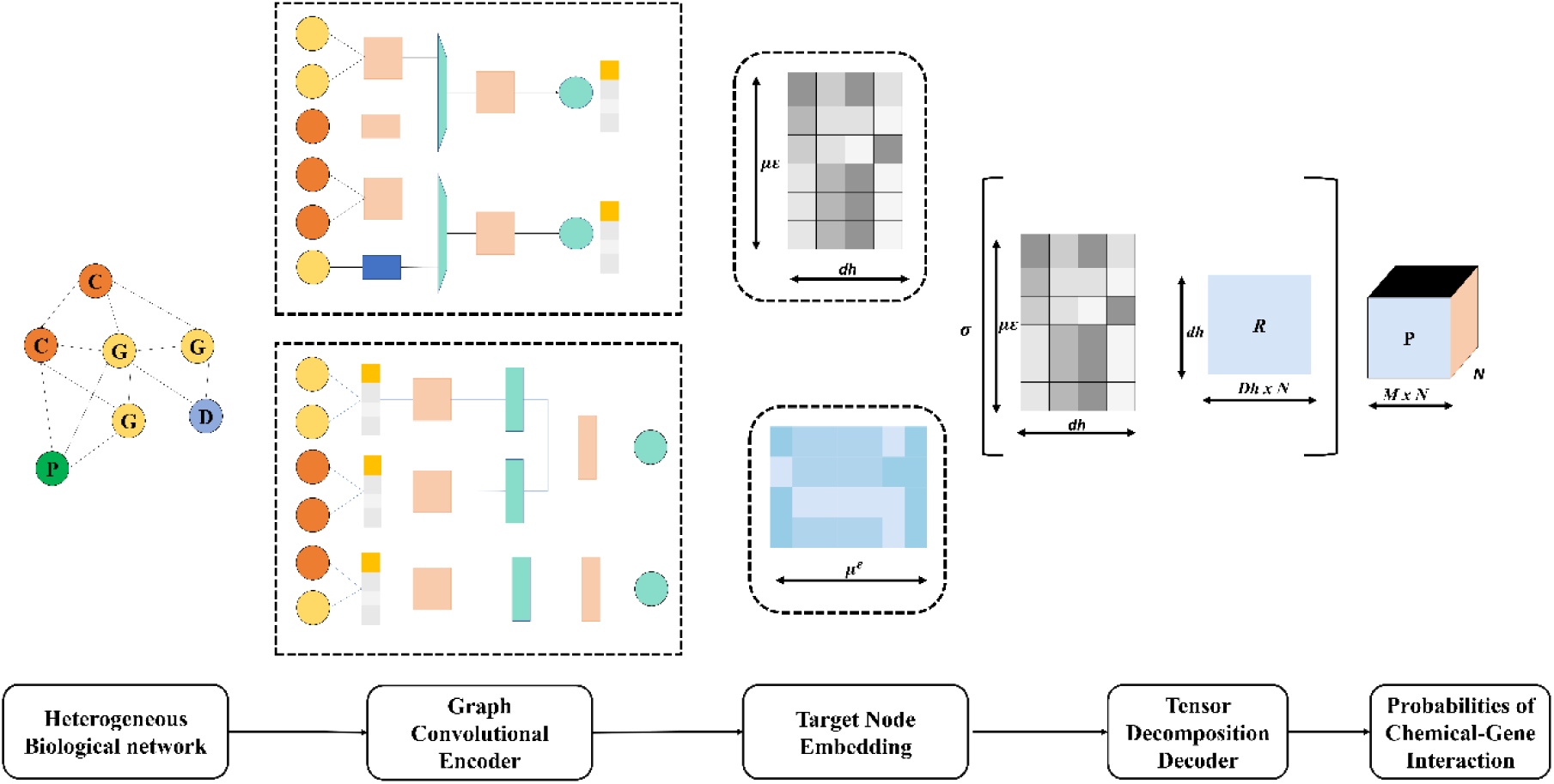
The overview of OMNI architecture - A pipeline for predicting CGIs from heterogeneous biological networks. A heterogeneous biological network with multiple node types, such as chemicals, genes, pathways, and diseases, and multi-relational edges is given as input. The relation-aware graph convolutional encoder performs per-relation message passing and produces compact embeddings for source/target nodes. The embeddings are organized into relation-specific latent tensors and passed to a tensor-decomposition decoder, which factorizes the latent tensor and produces normalized interaction likelihoods. Together, the encoder + tensor decoder convert complex, multi-relational network structures into low-dimensional representations and output probability scores for downstream CGIs prediction.

The details of the OMNI architecture are discussed in the following sections.

### Network construction

A total of seven subgraphs were considered in this work, including Gene-Gene (GG) subgraph, Chemical-Pathway (CP) subgraph, Gene-Pathway (GP) subgraph, Chemical-Disease (CD) subgraph, Gene-Disease (GD) subgraph, Chemical-Chemical (CC) subgraph and Chemical-Gene (CG) subgraph. These data sets were collected from various sources, including the SNAP Graph Library [21], the Comparative Toxicogenomics Database (CTD) [22] and the STITCH database [23]. The dataset statistics is depicted in **Table 1**.

**Table 1.** Statistics of the data set considered in this study.

|  | **Available** | **Duplicate** | **Unique** | **Source** |
| --- | --- | --- | --- | --- |
| Gene-Gene | 715612 | 0 | 715612 | SNAP Graph Library [21] |
| Chemical-Pathway | 1651193 | 0 | 1651193 | CTD [22] |
| Gene-Pathway | 135792 | 0 | 135792 | CTD [22] |
| Chemical-Disease | 9515673 | 6048639 | 3467034 | CTD [22] |
| Gene-Disease | 119200482 | 84927925 | 34272557 | CTD [22] |
| Chemical-Chemical | 17705799 | 11866621 | 5839178 | The STITCH database [23] |
| Chemical-Gene | 2983667 | 582747 | 2400920 | CTD [22] |

\*All the chemicals, genes, diseases and pathways are considered as nodes, and the interactions among themselves are considered as edges. These data created a final integrated multi-type interaction graph.

Many chemicals are used to treat various diseases by targeting specific genes in one or more pathways. Thus, in this study, diseases and pathways are considered as interaction entities, and the addition of this information can provide valuable insights in predicting CGIs. Thus, we constructed the GD-graph and CD-graph, which contain 34,272,557 and 3,467,034 interactions, respectively. We experimented with three combo-graphs 1. CG (relation between chemicals and genes), 2. CGP (relation between chemicals, genes and pathways) and 3. CGPD (relation between chemicals, genes, pathways and diseases) to investigate the performance of the OMNI model by adding disease as entities. The CGPD is seven times higher than CG interactions. The schematic representation of a partial heterogeneous graph is shown in **Figure 2**.

**Figure 2.**
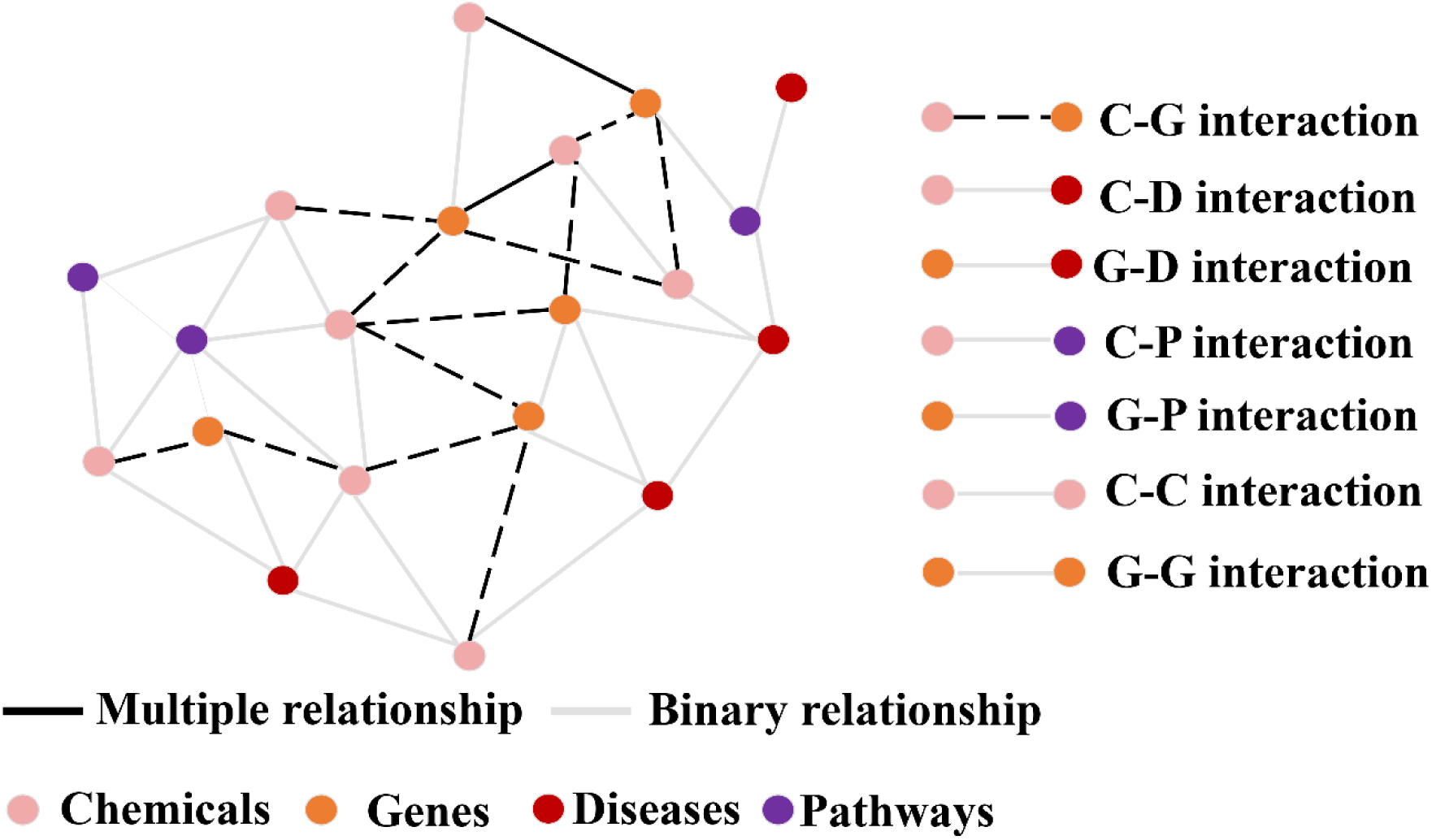
Illustration of the heterogeneous biological network used in OMNI, containing multiple node types (chemical, gene, disease, pathway) and diverse binary and multi-relational edges.

### Multi-view heterogeneous attention network encoder

Conventional graph encoders propagate and transform information across the network to capture complex relationships among nodes and edges. However, this method suffers from direct interactions, and interaction strength may weaken the effect of farther neighbours in the feature propagation process. Thus, in this study, we employed a novel architecture to model the heterogeneous graph from both local and global perspectives. The overall architecture of OMNI is represented in **Figure 3**.

**Figure 3.**
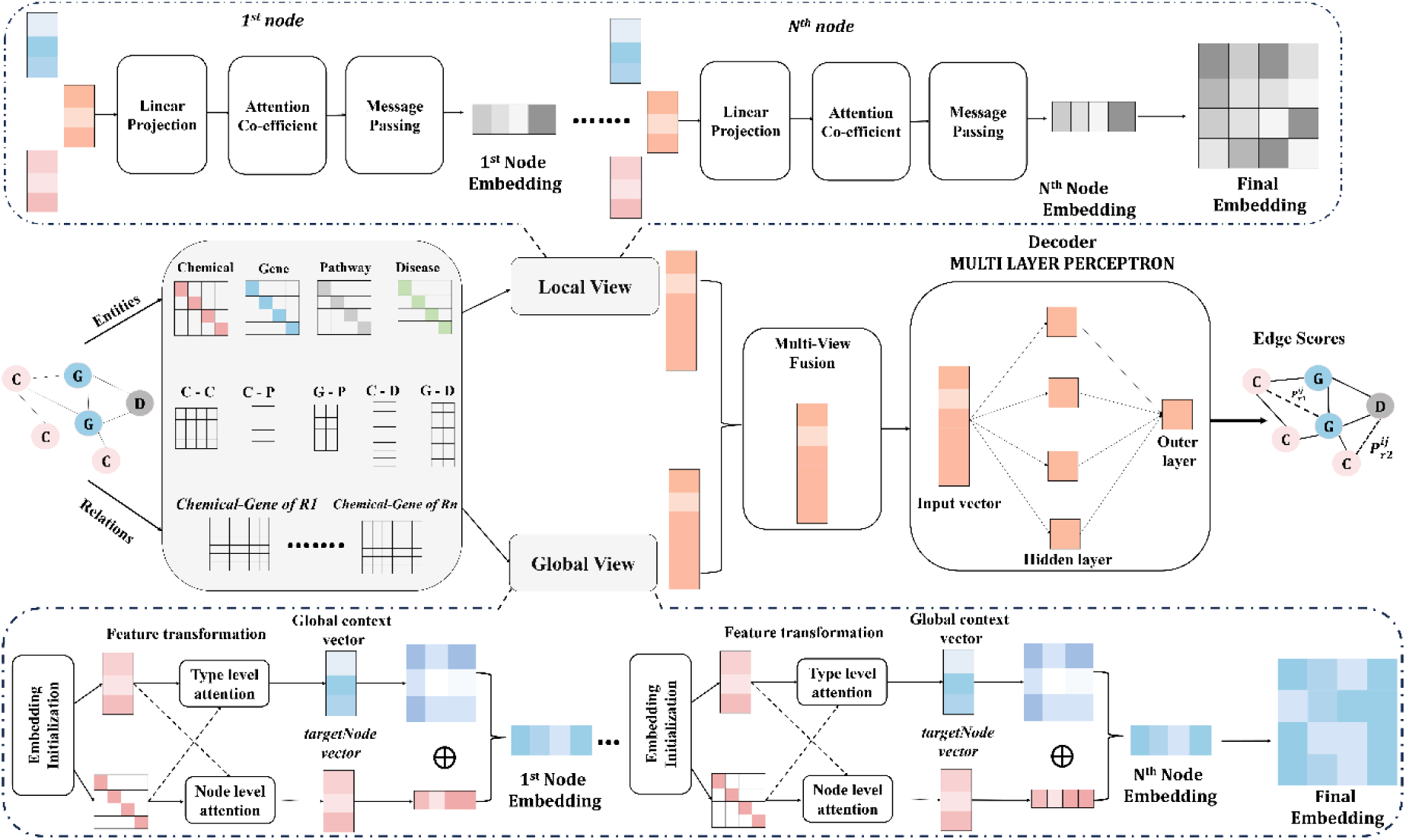
Overview of the OMNI architecture. The framework combines local and global relational views to learn expressive embeddings from a heterogeneous biological network. In the local view, nodes undergo linear projection, attention-based message passing, and refinement of their embeddings. In the global view, entity types and relations are encoded into adjacency matrices and enriched through type-level and node-level attention. The two views are employed multiple times to learn the local and global embeddings. Finally, the multi-view will fuse both the embeddings and pass them through an MLP decoder to produce final edge-level interaction scores.

### Feature transformation

In a heterogeneous graph, the different types of nodes (i.e., genes, diseases, pathways, chemicals) have distinct feature spaces in the heterogeneous space. Thus, a learnable type-aware feature transformation layer was applied to map them to the same feature space. Let *T* denote the set of node types, for each t ∈ 𝑇, let 𝑉_𝑡_be the set of nodes of type t, a learnable embedding 𝐸_𝑡_ ∈ ℝ^|𝑉𝑡|×𝑑^ and a linear projection 𝑊_𝑡_ ∈ ℝ^ℎ×𝑑^are used to encode node features into a common hidden space. *d* (feature_dim) denotes the vertex embedding width, h (hidden_dim) denotes the width after type projection, and K denotes the number of attention heads (local multi-head attention produces with *hK*).

For node 𝑣 ∈ 𝑉_𝑡_,

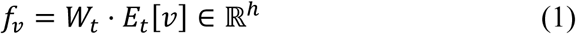

Here, t denotes the set of node types. A full-featured matrix 𝐹_𝑡_ = 𝑊_𝑡_𝐸_𝑡_ ∈ ℝ^|𝑉𝑡|×ℎ^ is cached for computational efficiency during global neighbor attention and is refreshed once per training epoch to avoid repeated full-graph projection computation.

### Local neighbour aggregation

The local neighbour information was aggregated similarly to simple HGCN [24], however, the difference lies in aggregating information from one-hop neighbours, unlike other popular methods such as BioNet. As discussed earlier, the target node representation is affected by weakly correlated neighbours and thus the farther node υ interaction may weaken the influence and be unable to capture the full information. Thus, we adopted a global view to aggregate the information of high-order neighbours.

The OMNI framework employs a local aggregation strategy rooted in relation-specific multi-head attention to model the fine-grained connectivity of nodes within heterogeneous graphs. This mechanism enables the encoder to selectively focus on relevant neighbours (within 1 hop neighbours) according to both edge type and node type, capturing subtle patterns in direct interactions. The model produces node features enriched with information from the most meaningful local structures by learning distinct attention weights for each relation.

### Linear projection

For each node *i* and each attention head *k*, OMNI uses a relation-specific learned projection to encode node features into an attention space. The feature vector 𝑓_𝑖_ of each node is transformed with a trainable weight matrix 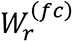 associated with the edge relation *r*, producing a head-specific feature:

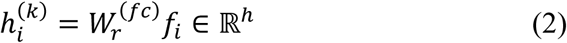

By generating relation- and attention head-specific embeddings, the model captures context-dependent variations, allowing precise modeling of heterogeneous interactions.

### Attention coefficients computation

The attention scores were further computed by the model for each source-destination neighbour pair *i*→*j* under relation *r* and head *k*. Using trainable vectors 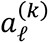 (for source) and 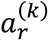 (for destination). The attention coefficient is calculated as:

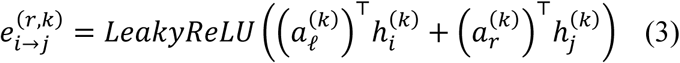

These raw scores are normalized using softmax for all incoming edges of each destination node *j*, yielding final attention coefficients:

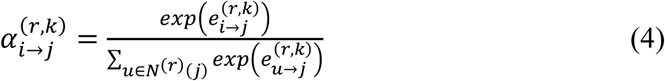

This process ensures that attention is distributed appropriately among neighbours, reflecting both relational and node-type information. 𝑁^(*r*)^(𝑗) is the set of 1-hop neighbours of *j* under relation r.

### Message Passing & Aggregation

Node representations are updated by gathering messages from neighbours, weighted by the computed attention scores. For each destination node *j:*

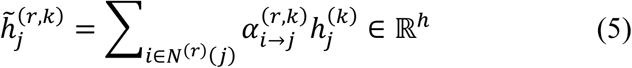

The outputs from all attention heads are concatenated:

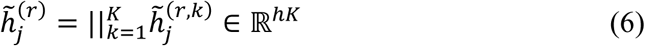

And then averaged over all relations ending in node-type *d:*

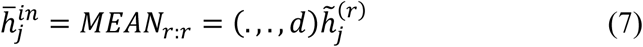

This step produces a summary embedding encapsulating both multi-head and multi-relation local information for each node.

### Residual, Normalization, and Activation

A residual connection incorporating the destination node’s own features (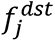) is added, expanded to match the attention head dimension to stabilize learning and preserve original node information:

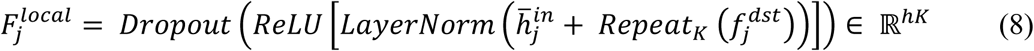

This normalization and activation pipeline ensures that aggregated features are scale-consistent and enriched, ready for subsequent fusion and global aggregation steps. The local view attention mechanism is represented in **Figure 4**.

**Figure 4.**
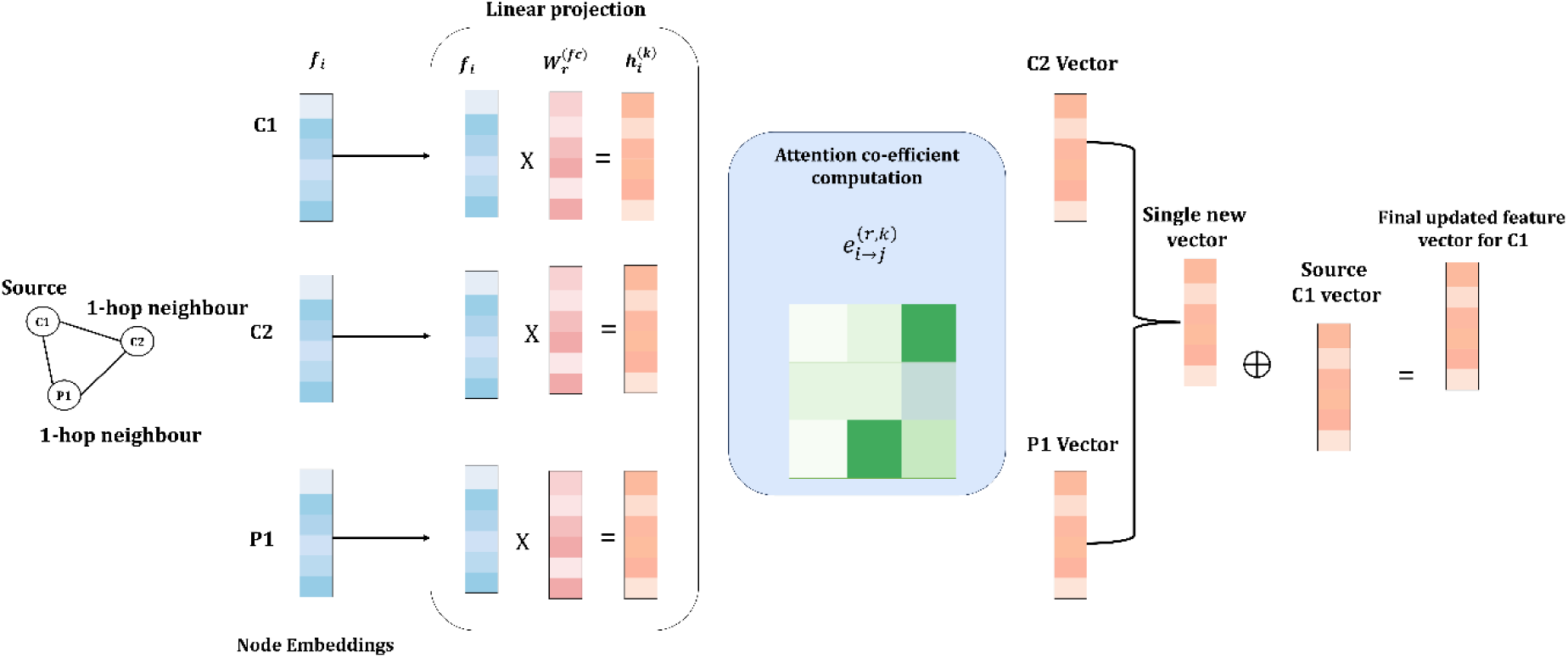
Local-view attention mechanism in OMNI. Node embeddings are first linearly projected, after which attention coefficients are computed across the 1-hop neighbourhood. Neighbour vectors are aggregated into a single message vector and combined with the source node’s embedding to produce the final updated feature representation.

### Global neighbour aggregation

The OMNI algorithm incorporates a global neighbour aggregation mechanism that captures high-order, distant interactions within the heterogeneous graph, thereby overcoming the limitations of direct, local neighbourhoods. This method effectively models the complex multi-type relationships that span multiple hops, which are common in biological networks. The global branch uses random walk sampling with restart (RWR) to collect a diverse set of heterogeneous neighbours for each node. It then employs a hierarchical attention mechanism: 1. Weighing the importance of different neighbor types, then 2. Assigning affinity scores to individual neighbors within those types. This two-level attention mechanism enables the model to selectively aggregate informative distant signals while filtering out less relevant nodes.

Finally, node features are aggregated with residual self-connections and linearly projected to align with the local view dimensions, thus enriching the node embeddings with comprehensive global context.

### Type attention score

The model assigns an attention score to each neighbour type 𝑡^′^ for a target node *v* to quantify the importance of information coming from that node type. This can be computed by

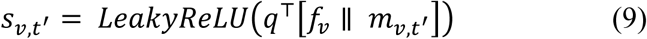

Where 𝑓_𝑣_is the transformed feature of node *v,* 𝑚_𝑣,𝑡_′ is a learned summary embedding for neighbors of type 𝑡^′^, and 𝑞 ∈ ℝ^2ℎ^ is a trainable query vector. These raw scores are normalized across all neighbor types using a softmax:

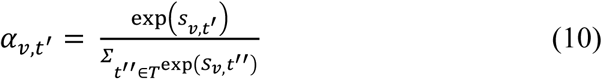

Assigning a normalized importance weight to each neighbour type.

### Node-level attention score

Within each neighbour type group, the importance of individual neighbours *u* is assessed via node-level attention:

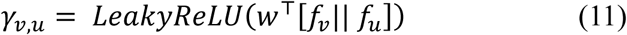

Where 𝑓_𝑢_ is the feature of neighbour *u,* and 𝑤 ∈ ℝ^2ℎ^ is a trainable affinity vector. The combined attention weight incorporates both node-level affinity and type-level weight:

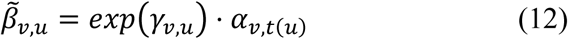

These weights are normalised over all global neighbours of *v*:

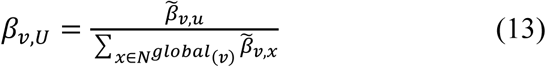

### Aggregation, Self-Connection and Projection

The final global representation for node *v* is computed as a weighted sum of neighbour features:

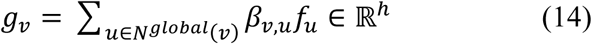

A residual self-connection adds the node’s own features:

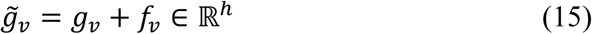

This combined vector is then linearly projected via a trainable weight matrix 𝑊^𝑔𝑙𝑜𝑏^ ∈ ℝ^ℎ𝐾×ℎ^ to match the dimensionality of the local multi-head attention output:

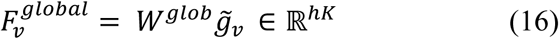

This projection allows seamless fusion of global and local views in subsequent processing steps.

This hierarchical attention-based global aggregation captures rich, heterogeneous, high-order dependencies, providing each node with broad contextual awareness complementary to local neighbourhood information. The global view attention mechanism is represented in **Figure 5**.

**Figure 5.**
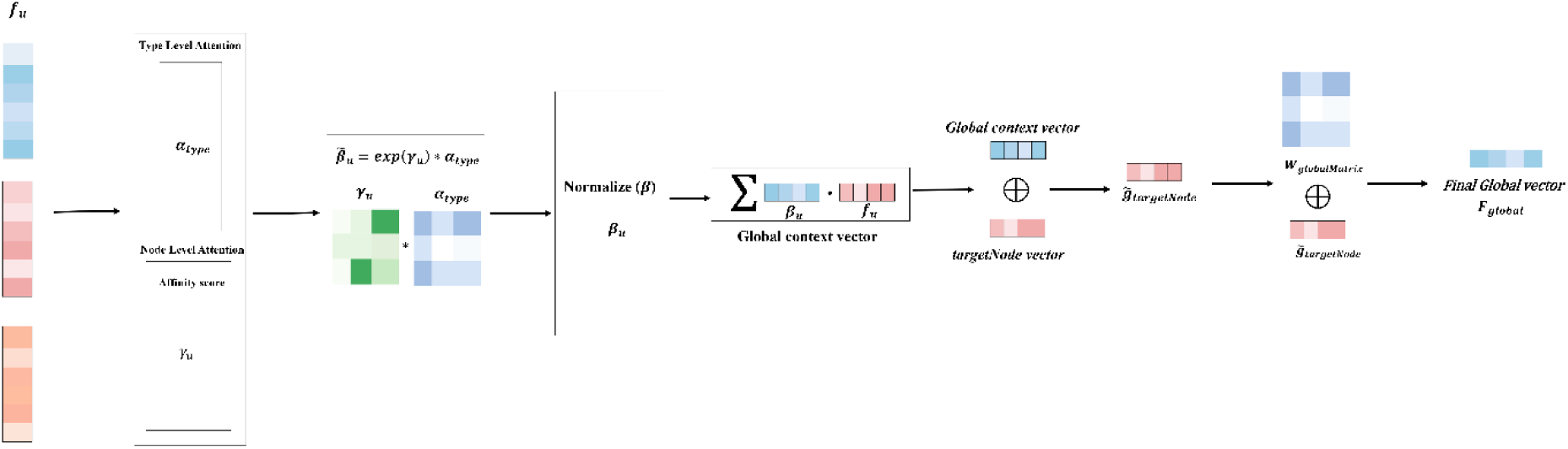
The global-view attention mechanism in OMNI. This module aggregates high-order relational information by first computing type-level and node-level attention scores across all entity types. The resulting affinity matrix is normalized and used to generate a global context vector. This vector is combined with the target node embedding through weighted fusion to produce the final global representation, capturing long-range dependencies beyond the local neighbourhood.

### Multi-view fusion

The OMNI model integrates node representations from the two views through a dedicated fusion mechanism, effectively combining complementary information captured from both local and global graph contexts. The local view, derived from relation-specific multi-head graph attention, encodes fine-grained and immediate neighborhood structure, while the global view aggregates high-order, multi-type neighbor information obtained through hierarchical attention. The model produces unified embeddings that reflect both short-range and long-range dependencies by concatenating these heterogeneous feature vectors and projecting them through a per-node-type linear transformation. This fusion step enhances representation richness and improves downstream edge prediction.

### Concatenation of local and global features

For each node *v* of type *t*, the outputs of the local neighbor aggregation 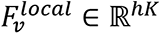 and the global neighbour aggregation 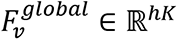 are concatenated to form a combined feature vector:

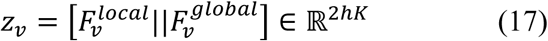

This concatenation preserves the distinct structural signals captured by each view while enabling their joint use in subsequent learning stages.

### Type-specific linear projection

To reconcile the concatenated vector into a final unified embedding, OMNI applies a learnable linear projection specific to each node type *t*:

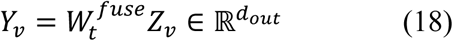

Where 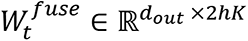 is a trainable projection matrix for type *t*, and 𝑑_𝑜𝑢𝑡_is the dimension of the fused representation. This per-type projection layer enables the model to adapt fusion mappings to the unique characteristics of different node types, improving representational alignment and expressiveness.

### Resulting unified node representation

The fused embeddings 𝑌_𝑣_serve as the final node-level representations that capture integrated multi-scale context from both local and global perspectives. These embeddings are subsequently used as input features for relation-specific decoders to predict edge likelihoods across heterogeneous relations, facilitating comprehensive and accurate graph completion. This multi-view fusion strategy leverages complementary modalities of graph context, balancing detailed local interactions with broad global structure to achieve state-of-the-art link prediction performance in complex heterogeneous networks.

### Relation-specific link decoding and training

In heterogeneous graph learning, predicting the existence and types of edges between nodes is crucial for tasks such as CGIs prediction. OMNI employs relation-specific decoders to estimate the likelihood of candidate edges for each canonical relation type defined in the heterogeneous graph. These decoders utilize the fused node representations as input, applying MLPs tailored to each relation to produce logits representing edge probabilities. The model trains these decoders via binary cross-entropy loss on observed positive edges and sampled negative edges, enabling effective discrimination of true interactions. This approach allows OMNI to simultaneously model multiple relation types while leveraging unified node embeddings, ensuring accurate and relation-aware edge inference.

### Relation-specific edge logit prediction

For each canonical relation *r* = (𝑠, 𝜌, 𝑑) where *s* and *d* are source and destination node types and 𝜌 is the relation type-OMNI predicts the likelihood of an edge between nodes *u* and *v* by concatenating their fused feature vectors:

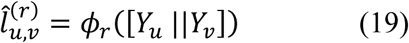

Here 𝑌_𝑢_ and 𝑌_𝑣_are the fused embeddings for nodes *u* and *v*, and 𝜙_*r*_ is a two-layer multi-layer perceptron (MLP) specific to relation *r.* This relation-specific MLP transforms concatenated node embeddings into a scalar logit representing the predicted edge strength or likelihood.

### Loss Function: Binary Cross-Entropy with Logits

OMNI trains the decoders using supervised learning with a binary cross-entropy loss applied to logits:

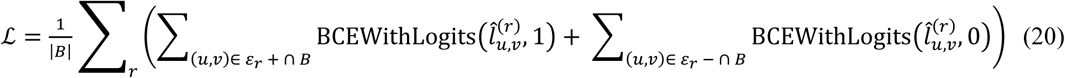

Where |𝐵| is the batch size 𝜀_*r*_ + and 𝜀_*r*_ − represent the sets of positive and sampled negative edges under relation *r*, respectively. This objective aims to achieve high prediction scores for true edges and low scores for negatives, thereby facilitating accurate edge classification.

### Optimization and Negative Sampling

Training utilizes the AdamW optimizer [25] to efficiently handle parameter updates, including weight decay regularization to prevent overfitting. Negative edges are uniformly sampled from the set of all non-observed edges per relation during training batches, maintaining a balanced dataset that aids robust edge discrimination. Model evaluation metrics, such as AUROC, AUPRC, Accuracy, and AP@K, are computed on the sigmoid-transformed output logits, providing a comprehensive performance assessment for link prediction. This relation-specific decoding and training scheme enables OMNI to perform multi-relational link prediction effectively across heterogeneous graphs, leveraging rich, fused node embeddings while preserving relation-dependent predictive nuance.

## Results

### Experimental settings

All computational calculations were performed on the CSIR - Fourth Paradigm Institute (CSIR–4PI) supercomputing facility utilizing NVIDIA A100-SXM4 80GB GPUs. DL frameworks Pytorch and *Pytorch Lightning* were adopted to generate the OMNI code. The binary cross-entropy with logits was considered as the loss function, and it is optimized using the AdamW optimizer. The parameters used in the model are depicted in **Table 2**.

**Table 2.** The parameters used in OMNI.

| **Parameter** | **Description** | **Value** |
| --- | --- | --- |
| epochs | The number of training epochs | 30 |
| batch_size | The number of samples per training step | 4096 |
| feature_dim | The initial node feature/embedding dimension | 128 |
| hidden_dim | The hidden layer dimension in the encoder | 64 |
| out_dim | The output embedding dimension | 64 |
| num_heads | The number of attention heads in GAT layers | 4 |
| dropout | The dropout rate for regularization | 0.3 |
| lr | The learning rate of the AdamW optimizer | 0.0001 |
| weight_decay | The weight decay for AdamW optimizer | 0.00001 |
| fanouts | The neighbor sampling fanouts for each GNN layer | 15,10 |
| neg_k | The number of negative samples per positive sample | 1 |
| rwr_len | The length of random walk paths for global attention | 10 |
| rwr_restart_prob | The restart probability in Random Walk with Restart | 0.1 |
| rwr_top_k | The top-k neighbors to keep from RWR for global context | 20 |

### Performance evaluation

Three different classical evaluation metrics were calculated to comprehensively evaluate the performance of OMNI, including area under the receiver operating characteristic curve (AUROC), accuracy, and area under the precision-recall (PR) curve (AUPRC). Initially, all the CGI instances were randomly divided into a training set, a test set and a validation set with a ratio of 8:1:1 per interaction type. The performance of OMNI has been compared with that of other state-of-the-art methods and the BioNet model on three different datasets (i.e., CG, CGP and CGPD).

The performance of OMNI is compared with different baseline algorithms (i.e. CGINet [26], Node2Vec [27]) and one more advanced method BioNet. As illustrated in **Table 3**, the OMNI method consistently outperformed the other methods in all three different metrics, including AUROC, AUPRC, and accuracy. To note, the Node2Vec algorithm could not effectively handle the larger dataset (CGPD), thus, they were evaluated only on the CG and CGP datasets. The experimental results clearly indicate that the OMNI outperforms the different methods with the same subgraphs. This is mainly due to two reasons: 1. The adopted OMNI method models the heterogeneous graph from both local and global views, which helps to capture the effect of farther nodes effectively 2. OMNI adopts GPU parallelization using *PyTorch Lightning* and multi-node job distribution. This enormously helps to optimize the training process, where it trains each relation very effectively in a reasonable time period.

**Table 3.** Performance comparison of OMNI models with other baseline approaches considered in this study.

| **Model** | **Dataset** | **AUROC** | **AUPRC** | **Accuracy** | **AP@20** |
| --- | --- | --- | --- | --- | --- |
| BioNet | CG | 0.8212 | 0.8506 | 0.5045 | 1.0000 |
|  | CGP | 0.8373 | 0.8512 | 0.5146 | 1.0000 |
|  | CGPD | 0.8298 | 0.8539 | 0.5027 | 1.0000 |
| CGINet | CG | 0.7820 | 0.7993 | 0.6181 | 1.0000 |
|  | CGP | 0.7594 | 0.7827 | 0.7050 | 0.3940 |
|  | CGPD | 0.7881 | 0.7974 | 0.6426 | 0.8033 |
| Node2Vec | CG | 0.8150 | 0.7833 | 0.7100 | 0.0000 |
|  | CGP | 0.8333 | 0.7417 | 0.8007 | 0.0000 |
|  | CGPD | - | - | - | - |
| OMNI | CG | 0.8800 | 0.8593 | 0.8119 | 1.0000 |
|  | CGP | 0.8910 | 0.8709 | 0.8229 | 1.0000 |
|  | CGPD | 0.9089 | 0.8894 | 0.8415 | 1.0000 |

Additionally, we benchmarked different decoders (i.e. Multi Layer Perceptron (MLP) [16], MultX [17], TransE [18], Reverse [19]) methods and the MLP performance is relatively better than the other decoders (**Table 4**). Additionally, it is worth noting that introducing a new set of nodes and edges can improve the performance of OMNI. Thus, OMNI with CGPD yields better results when compared to all other methods.

**Table 4.** Comparison of different decoders applied in the OMNI models.

| **Model** | **Dataset** | **AUROC** | **AUPRC** | **Accuracy** | **AP@20** |
| --- | --- | --- | --- | --- | --- |
| DistMult | Train | 0.9055 | 0.8711 | 0.8471 | 1.0000 |
|  | Val | 0.9010 | 0.8801 | 0.8336 | 1.0000 |
|  | Test | 0.8875 | 0.8667 | 0.8209 | 1.0000 |
| ComplEX | Train | 0.9064 | 0.8726 | 0.8480 | 1.0000 |
|  | Val | 0.8984 | 0.8765 | 0.8314 | 0.9551 |
|  | Test | 0.8841 | 0.8638 | 0.8173 | 0.9971 |
| RESCAL | Train | 0.9058 | 0.8716 | 0.8473 | 1.0000 |
|  | Val | 0.8983 | 0.8768 | 0.8314 | 1.0000 |
|  | Test | 0.8858 | 0.8654 | 0.8197 | 1.0000 |
| MLP | Train | 0.9097 | 0.8773 | 0.8516 | 1.0000 |
|  | Val | 0.9089 | 0.8894 | 0.8415 | 1.0000 |
|  | Test | 0.8862 | 0.8724 | 0.8215 | 1.0000 |

\*All the calculations are done with the CGPD dataset. The MLP performance is relatively better than the other decoders.

### Computational optimization

The device memory of a single NVIDIA A100 GPU (80GB) is insufficient to accommodate the full OMNI-CGPD model and its large-scale heterogeneous graph data. During single-device experiments, the model rapidly consumed more than 80GB of GPU memory. Thus, we adopted distributed data-parallel training across four NVIDIA A100 (80GB) GPUs, enabling OMNI-CGPD to process the entire biomedical knowledge graph without memory overflow and with significantly improved training efficiency. We evaluated the parallel performance of OMNI using multi-GPU settings. **Figure 6** illustrates the training time per epoch for OMNI-CG, OMNI-CGP and OMNI-CGPD using different numbers of GPUs.

**Figure 6.**
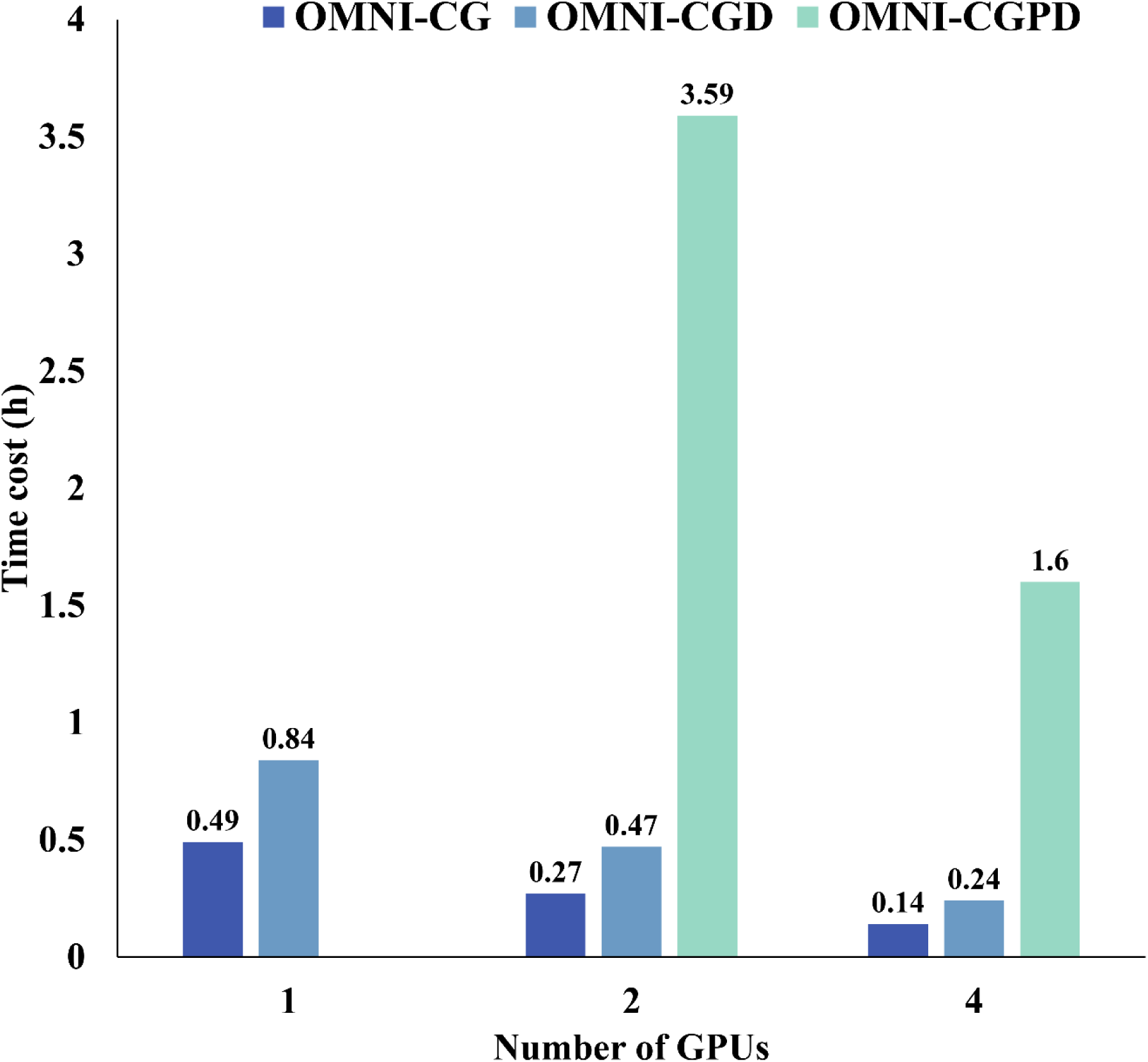
Comparison of the training time for OMNI variants (OMNI-CG, OMNI-CGD, OMNI-CGPD) across different numbers of GPUs. Each bar shows the average time cost (in hours), illustrating how model complexity and GPU count affect computational efficiency.

In contrast, the full-scale OMNI-CGPD model requires distributed computation. A single GPU cannot fulfil the computation of OMNI-CGPD, whereas with four GPUs operating under the Distributed Data Parallel (DDP) strategy, the per-epoch time decreases to 1.6 hours. The corresponding speedup and parallel efficiency are computed as follows:

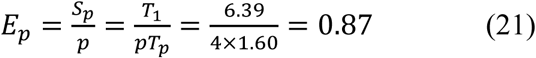

Where E_p_ represent parallel efficiency, S_p_ represents speedup, T_1_ refers to the execution time of the sequential implementation, T_p_ represents the execution time of the parallel implementation, and *p* indicates the number of GPUs.

This configuration achieved a speedup of 3.48x and a parallel efficiency of 87%, effectively reducing the training time per epoch by nearly fourfold. Each GPU utilized approximately 77GB of its available 80GB memory, demonstrating that the full model computation can only be realized through distributed GPU memory. These results confirm that multi-GPU parallelism substantially accelerated OMNI-CGPD training while maintaining full-scale model capacity and stable convergence.

### Ablation study

We conducted an ablation study comparing three model configurations: (i) Only global view, (ii) Only local view, and (iii) the combined model (Local + Global) to examine the contribution of each architectural component. All variants were trained and evaluated under identical conditions for a single epoch, and performance was assessed across multiple metrics, including AUROC, AUPRC, Accuracy, and AP@20 on the train, validation, and test sets. The results are discussed below,

The global view model demonstrated moderate performance, achieving a Test AUROC of 0.5908, AUPRC of 0.6009, and accuracy of 0.5569. These values indicate that while the global-level representation captures broad interaction patterns, it lacks fine-grained discriminatory power. The consistently perfect AP@20 suggests strong ranking ability for the topmost predictions, but the overall classification capability remains limited.

Introducing local structural information led to a substantial performance gain across all evaluation metrics. The local view model achieved a test AUROC of 0.8764, AUPRC of 0.8561, and accuracy of 0.8082, showing that fine-grained contextual details play a crucial role in distinguishing biologically relevant interactions. Compared to the global view model, AUROC improved by nearly 0.29, and similar improvements were observed in AUPRC and Accuracy. This confirms that localized representation learning is highly informative for the task.

Integrating both global and local views, the combined model achieved the best overall performance across all datasets and metrics, demonstrating the complementary nature of the two representations. On the test set, the model obtained the highest AUROC (0.8774), AUPRC (0.8583), and accuracy (0.8092), surpassing both ablated versions. Although the improvements are incremental compared to the local view model, the combination of global and local views provides meaningful context that enhances predictive consistency and refinement. Notably, the model maintains a good AP@20 score across all phases, reinforcing its robustness when prioritising high-confidence predictions. Overall, across all metrics and datasets, the combined model consistently delivers the strongest performance, demonstrating that joint encoding of global interactions and local fine-grained information yields the most complete and discriminative representation. The ablated results clearly show that while local features drive most of the predictive strength, incorporating global structure provides additional contextual reinforcement, leading to the highest overall accuracy and ranking quality. The ablation study results are depicted in **Table 5**.

**Table 5.**
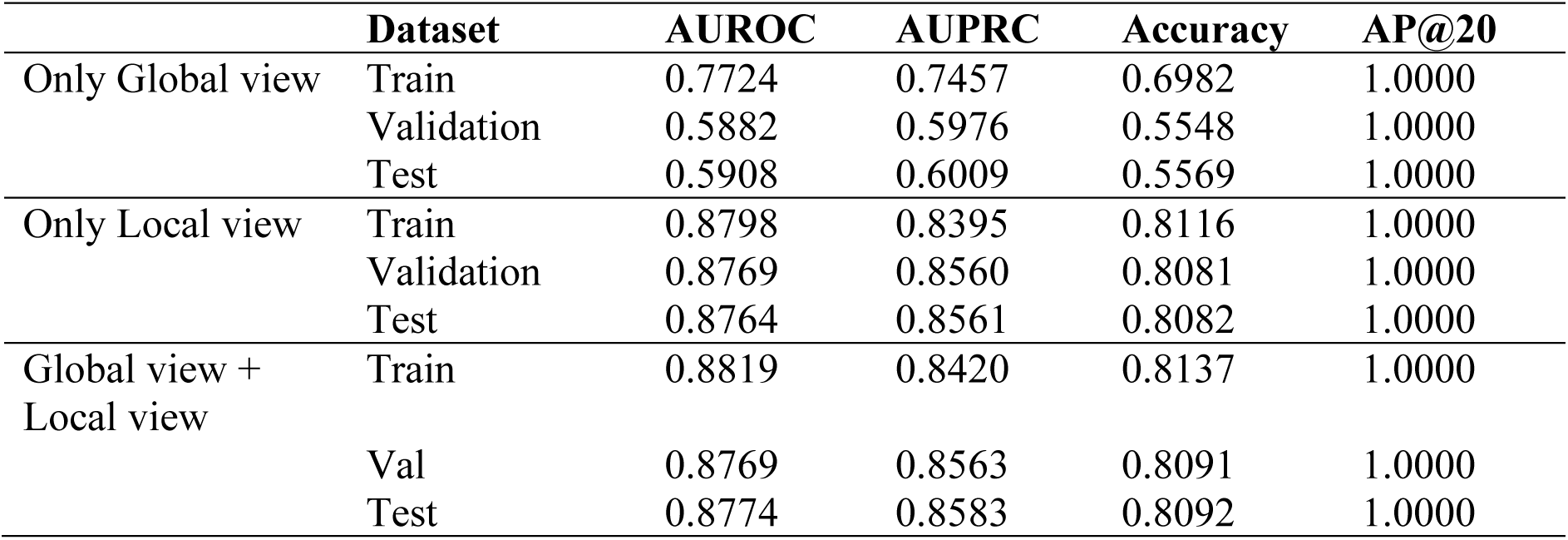
Ablation study results.

### Case study

Several case studies were conducted to further explore the applicability and usage of OMNI in enhancing biomedical studies.

### Identification of cancer targets and small molecule pairs

#### Case study I

There is considerable effort in cancer research aimed at various pathogenic mechanisms and modulating these pathogenic sites. In a pan-cancer proteogenomics study [28], 1043 patients across 10 cancer types and 2,863 druggable proteins were analysed. In this study, the authors have collected drug target information from various resources, including DrugBank [29], Guide to Pharmacology (GtoPdb) [30], the Drug Gene Interaction Database [31], and the *in silico* surfaceome [32]. The data is classified into five tiers. We considered Tier 3 from this study, which contains 448 targets inhibited by drugs from investigational/experimental, especially epigenetic drugs. The collected data was compared with the OMNI dataset, and the existing relation pairs were removed from the source database. The remaining predictions were ranked in descending order and listed in **Table S1**. A high score indicates a higher predicted probability of the potential interaction. Furthermore, we evaluate these interactions by searching for entity pairs in various sources, including PubMed and the Google Scholar search engine, to provide supportive evidence in the literature. A list of partially verifiable CGIs related to cancer that are predicted using OMNI is depicted in **Table 5**, along with the evidence. The table represents the lists of chemical (node c) and gene (node v) pairs, along with possible relation types and literature evidence. The score 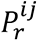 represents the predicted probability of the link 𝑒_𝑖𝑗_ = (𝑣_𝑖_, *r*, 𝑣_𝑗_) generated by the OMNI tensor factorization decoder.

The WYE-354 is an ATP-competitive mTOR kinase inhibitor that blocks both mTORC1 and mTORC2 signalling by reducing the phosphorylation of downstream substrates, such as p-S6K(T389) and p-AKT(S473), and producing dose-dependent suppression of mTOR activity and cell proliferation in preclinical models. The A-674563 molecule is a selective AKT1 inhibitor that reduces AKT kinase signalling by decreasing the phosphorylation of downstream AKT substrates, thereby slowing tumour cell proliferation by decreasing AKT1 activity. The JNK-IN-8 is a covalent/irreversible JNK family inhibitor that suppresses JNK phosphorylation activity and reduces phosphorylation of the direct substrate c-Jun in cells, thereby decreasing MAPK8/JNK signalling and downstream inflammatory responses. The clascoterone is an androgen receptor inhibitor that competes with androgens (i.e., DHT), with clinical and *in vitro* data supporting its decrease in androgenic activity. The molecule Pamapimod is a selective small-molecule inhibitor of p38α MAP kinase, inhibiting p38 enzymatic activity and downstream inflammatory cytokine production in both preclinical and clinical studies. Adapalene modulates the AP-1/c-Jun related transcriptional pathway in keratinocytes and inflammatory skin models. The adapalene downregulates AP-1-driven inflammatory signalling and thereby reduces c-Jun-dependent expression in treated skin. Docetaxel affects the stability of MAPT (tau protein) by promoting tubulin polymerization. Docetaxel has an effect on microtubule dynamics and interacts functionally with microtubule-associated proteins such as MAPT/tau and reduces the stability of microtubules. The Oxaprozin decreases the PTGS1 activity. This molecule is a propionic acid NSAID that inhibits cyclooxygenase enzymes (COX-1 and COX-2 also known as PTGS1 and PTGS2) and reduces prostaglandin synthesis, thereby decreasing COX enzymatic activity *in vivo*. Afatinib is an irreversible covalent ErbB family tyrosine kinase inhibitor that blocks the kinase activity of EGFR and ErbB2 (HER2) and reduces receptor autophosphorylation and downstream signalling. The Auranofin decreases the activity by inhibiting IκB kinase (IKK) activity and thereby suppresses NF-kB signalling in biochemical and cell experiments and produces anti-inflammatory effects. The Brompheniramine is a classical H1-antihistamine that acts as an H1 receptor antagonist/inverse agonist (competes for histamine binding), thereby reducing histamine-driven H1 receptor activity and downstream responses. Caffeine is a non-selective adenosine receptor antagonist (A1, A2A, A2B and A3) and thus functionally reduces signalling through adenosine receptors, including A3 (by antagonism/inverse agonism), producing decreased ADORA3-mediated activity. Cyclizine is a first-generation antihistamine that binds to H1 receptors, preventing histamine binding (antagonistic activity) and decreasing histamine–HRH1 interactions and functional H1 signalling. Diethylpropion is an amphetamine-like anorectic that increases synaptic dopamine levels primarily *via* promoting dopamine release and/or interfering with dopamine reuptake. Diflunisal is a salicylate derivative NSAID that inhibits cyclooxygenase enzymes (COX-1 and COX-2 / PTGS1 and PTGS2), decreasing COX-2 catalytic activity and prostaglandin production and mediating anti-inflammatory/analgesic effects.

#### Case study II

Additionally, a list of top genes associated with cancer was retrieved from various sources. The OMNI was used to obtain all related chemical predictions and exclude existing relation pairs from the source database. The remaining predictions were ranked in descending order and listed in **Table S2**. The high potential interaction is represented by a high score. In the next step, we searched for literature evidence supporting the predicted relation in various resources, including Google Scholar and PubMed, to find corroborating evidence. **Table 6** illustrates the predicted partially verifiable relations.

**Table 6.** The predicted CGIs of cancer predicted by OMNI.

| Chemical-gene pairs | Probability<br>score | Relation type | Evidence |
| --- | --- | --- | --- |
| WYE-354, MTOR | 0.9906 | decreases^activity | [33] |
| A-674563, AKT1 | 0.9968 | decreases^activity | [34] |
| JNK-IN-8, MPK8 | 0.9942 | decreases^activity | [35] |
| Clascoterone, AR | 0.9982 | decreases^activity | [36] |
| Pamapimod, MAPK14 | 0.9959 | decreases^activity | [37] |
| Oxaprozin, PTGS1 | 0.9970 | decreases^activity | [38] |
| Afatinib, ERBB2 | 0.9900 | decreases^activity | [39] |
| Auranofin, IKBKB | 0.9833 | decreases^activity | [40] |
| Brompheniramine,<br>HRH1 | 0.9992 | decreases^activity | [41] |
| Caffeine, ADORA3 | 0.090378 | decreases^activity | [42] |
| Diflunisal, PTGS2 | 0.9977 | decreases^activity | [43] |

The activity of Atorvastatin with CAMKK2 was proven by several preclinical studies. Patients treated with atorvastatin have increased CaMKK activity in endothelial and metabolic models. Multiple studies have shown that Withaferin A induces the accumulation and increased expression/activation of p53, leading to p53-dependent apoptosis in cancer cells. Garcinol is an epigenetic modifier that has been reported to induce p53 expression and increase p53 acetylation by promoting p53 accumulation and pro-apoptotic functions in various *in vitro* studies. Lidocaine decreases vimentin expression as reported in several cell studies and inhibits migration/EMT markers in cancer and epithelial models (i.e., lidocaine decreased VIM expression in certain experimental contexts). Doxycycline has been shown to reduce production of PDGF ligands and to suppress PDGF/PDGFR signalling in cellular fibrosis and tumor models (reduced PDGF-AA production and inhibition of PDGF/PDGFR signalling/phosphorylation), consistent with an overall reduction in PDGFR pathway expression/signalling in these contexts. Guadecitabine, a next-generation DNMT inhibitor, causes DNMT1 depletion in cancer cells (reduced DNMT1 protein abundance) and hypomethylation of target loci in multiple preclinical and clinical studies. Capmatinib is a potent, selective MET tyrosine kinase inhibitor that blocks MET kinase activity and has demonstrated *in vitro* and *in vivo* activity against MET-driven tumors, including NSCLC. The Eflornithine is an irreversible inhibitor of ornithine decarboxylase (ODC1) that reduces ODC enzymatic activity and polyamine synthesis in tissues, with strong evidence for decreased ODC1 activity. Entrectinib is an oral small-molecule inhibitor targeting TRK, ROS1 and ALK kinases, and it potently inhibits ROS1 kinase activity and produces clinical responses in ROS1-fusion positive NSCLC. Nintedanib is a small-molecule tyrosine kinase inhibitor that targets VEGFR by inhibiting receptor tyrosine kinase activation and downstream pro-angiogenic/pro-fibrotic signalling, with clinical data supporting its role in decreasing VEGFR activity.

#### Case study III

##### OMNI in deciphering phytochemicals gene interactions

The usability of OMNI is further evaluated by exploring the gene-phytochemicals interactions. The VDR is considered for this analysis as it is crucial for several physiological functions. As a first step, we considered all the chemicals available in the network and omitted those with a direct edge to the VDR. The probability scores for all interaction types were calculated for all remaining chemicals and listed in the *Supplementary Information*. We further evaluated the relationship between the VDR and the phytochemicals, along with partially verifiable interactions. The results are listed in **Table 7**. Interestingly, all the chemicals in the selected list have a good probability score with the VDR. Further, the phytochemicals and their proven VDR mechanisms are discussed in detail below,

**Table 7.**
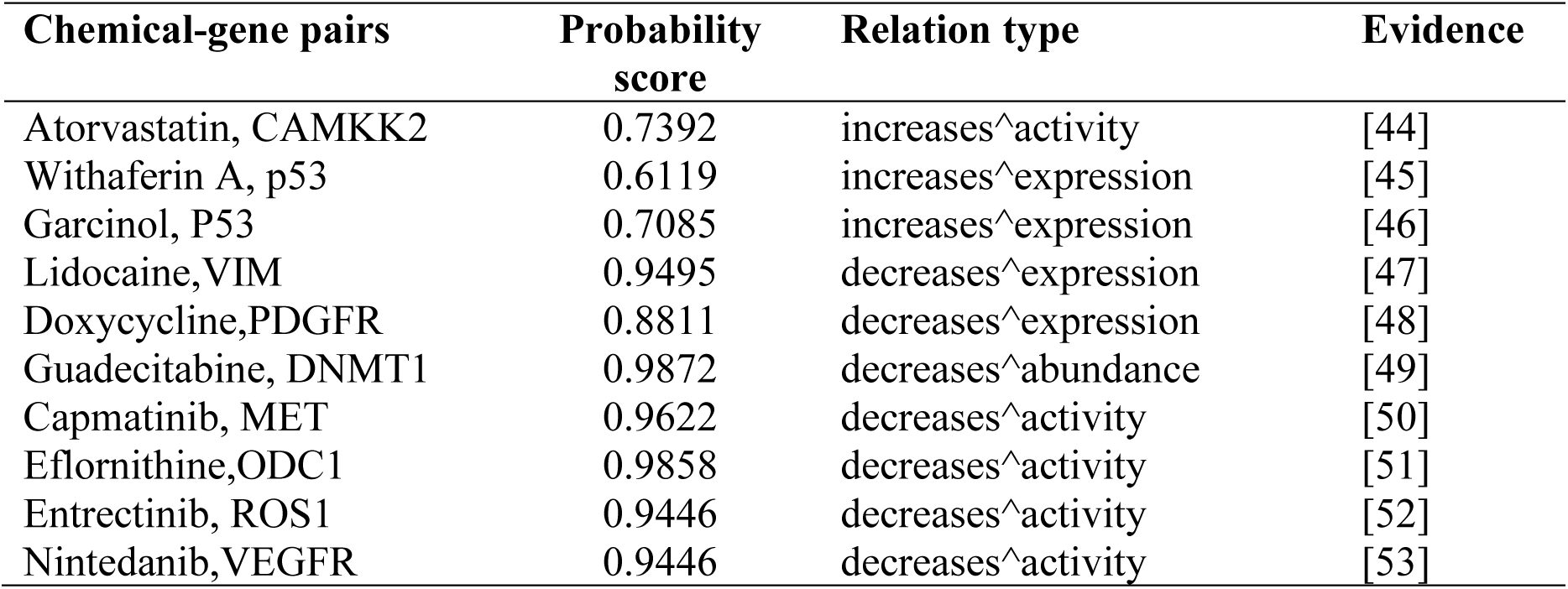
The predicted CGIs of cancer predicted by OMNI.

**Table 8.** Phytochemicals – VDR interaction predicted by OMNI.

| Phytochemicals | Probability score | Relation type | Evidence |
| --- | --- | --- | --- |
| Astragalus polysaccharide | 0.9794 | affects^abundance | [54] |
|  | 0.9957 | increases^activity |  |
| Bufalin | 0.9899 | increases^activity | [54] |
|  | 0.9980 | increases^localization |  |
|  | 0.9893 | affects^binding |  |
|  | 0.9815 | affects^response_to_substance |  |
| Tanshinone IIA | 0.9654 | increases^expression | [55] |
|  | 0.9882 | increases^activity |  |
|  | 0.9937 | affects^response_to_substance |  |
| Vitexin | 0.9852 | increases^activity | [56] |
|  | 0.9888 | affects^binding |  |
| Berberine | 0.9789 | increases^expression | [57] |
|  | 0.9908 | increases^activity |  |
|  | 0.9895 | affects^response_to_substance |  |
| Astragaloside | 0.9888 | increases^expression | [58] |
|  | 0.9908 | increases^activity |  |
| Parthenolide | 0.9915 | increases^expression | [59] |
|  | 0.9891 | affects^activity |  |

The phytochemical Astragalus polysaccharide (APS) exhibits multiple VDR regulatory effects, and experimental studies have shown that APS can modulate Vitamin D metabolic enzymes (i.e., CYP27B1 and CYP24A1) and alter the VDR abundance. In septic or adrenal models, APS slightly decreases VDR expression. However, in other contexts (e.g., immune modulation and gut microbiota regulation), it enhances VDR signalling, improving inflammatory control and metabolic balance. The other bioactive compound, Bufalin from *Toad venom (Chansu),* has known functional connections to VDR. It increases VDR activity by potentiating the transcriptional effects of 1,25-dihydroxyvitamin D₃ on canonical target genes such as *CYP24A1* and *cathelicidin (CAMP)*. The other phytochemical, Tanshinone IIA, is a diterpene quinone from *Salvia miltiorrhiza (Danshen)* that can act as a VDR modulator. This molecule upregulates VDR expression and reactivates VDR/Wnt/β-catenin pathways, leading to suppression of epithelial–mesenchymal transition (EMT) and fibrotic progression in kidney and liver fibrosis models. Vitexin is identified as a novel VDR agonist, which is a flavone glycoside found in *Vitex* species and other medicinal plants. It directly binds to the ligand-binding domain of VDR and activates its transcriptional activity, as shown from computational studies and biochemical assays. Another phytochemical, Berberine, is an isoquinoline alkaloid derived from *Berberis* species, which increases both VDR expression and transcriptional activity. It enhances VDR-mediated barrier-protective effects in intestinal epithelial cells and upregulates VDR mRNA and protein in gut and adipose tissue. The Astragaloside is a key compound from *Astragalus membranaceus* that upregulates the VDR expression, especially in cardiac and renal tissues under stress. It activates the VDR-related gene network, including regulators of vitamin D metabolism (CYP27B1 and CYP24A1). Parthenolide, a sesquiterpene lactone from *Tanacetum parthenium* (known as feverfew) has a few direct evidence but is predicted to enhance VDR expression and modulate its activity indirectly. Its anti-inflammatory actions, primarily through NF-κB inhibition, intersect with VDR pathways, as VDR is known to antagonize NF-κB-mediated transcription.

## Discussion

In the data science era, the abundance of gene-related and drug-related data presents tremendous opportunities for developing new insights and improved approaches to drug discovery. The fusion of heterogeneous biomedical data from various sources can provide a systematic understanding of biopharmaceutical mechanisms, offering more comprehensive and effective support for drug repurposing and increasing the accuracy of predictions. Other than CGIs, various types of relations, including ∼omics data, side effects data, etc., are essential for drug repurposing. Recent studies have garnered significant attention by exemplifying various types of interactions other than CGI. These can be broadly classified as target-centred models and disease-centred models.

NeoDTI [61] is an end-to-end learning model that integrates diverse information from heterogeneous network data and learn topology-preserving representations of targets and drugs to facilitate DTI prediction. In another study, a network-based computational framework, known as an arbitrary-order proximity embedded deep forest approach, was developed for predicting DTIs. This method learns a low-dimensional vector representation from a heterogeneous biological network that connects drugs, targets and diseases. Different types of information, including chemical, genomic, phenotypic, and network profiles, were considered for constructing the network, which encompasses drugs, diseases, and proteins. This method outperformed different state-of-the-art methods. Mohamed et. al. developed TriModel [62], a specific knowledge graph embedding model that connects targets and drugs. This is primarily derived from knowledge graphs generated using biological entities related to drugs and targets available from different biological databases, including UniProt [63], KEGG [64] and DrugBank [29]. This method outperformed other methods in terms of ROC and precision-recall curves.

There are a few other studies that focus on the disease-centred models. Wang et. al. [65] developed a bipartite GCN using drug-protein, disease-protein and protein-protein interaction pairs. They proposed BiFusion, a bipartite GCN for drug repurposing through heterogeneous information fusion. In another study [66], a group of scientists constructed a knowledge graph with 15 million edges across 39 different types of relationships connecting drugs, diseases, genes/proteins, pathways and expression data. A network-based DL method was applied to identify 41 repurposable drugs using Amazon’s AWS computing resources. deepDR [67] is another network-based DL approach for *in silico* drug repurposing by integrating 10 different networks including drug, side-effects, and targets. The deepDR effectively learns high-level features of frugs from heterogeneous networks by a multi-modal deep autoencoder.

Recent advances, such as NeoDTI, TriModel, and deepDR, have underscored the importance of integrating heterogeneous biological information for predicting drug–target interaction (DTI) and drug repurposing. These models successfully learn latent representations from complex biological networks. However, most models treat heterogeneous sources as a single unified structure, thereby neglecting view-specific dependencies among biological entities. For example, NeoDTI captures topological relations but lacks explicit multi-view representation, while TriModel focuses on knowledge graph embeddings without preserving the broader network topology. Similarly, deepDR employs a multimodal autoencoder to integrate multiple biological networks, yet this approach tends to compress network-specific information, potentially limiting the ability to capture higher-order associations across biological domains.

In contrast, our proposed multi-view heterogeneous GCN framework integrates both local and global perspectives of the biological network to derive more informative and topology-consistent representations. The local view retains immediate relational patterns, such as drug–target and protein–protein interactions, while the global view captures higher-order dependencies that span across the heterogeneous network. This joint learning strategy enables the effective fusion of information across multiple biological contexts, enhancing the capacity to model complex molecular mechanisms. Comparative evaluations demonstrate that our framework consistently outperforms existing methods, particularly in identifying novel or weakly connected drug–target pairs. By coupling multi-view learning with heterogeneous graph convolution, the proposed approach provides a robust and biologically interpretable foundation for in silico drug discovery and repurposing.

Beyond predictive performance, the model’s architecture facilitates biological interpretability by linking learned representations to specific molecular and pathway-level features. The separation of local and global network views enables tracing of predicted interactions back to relevant subnetworks, such as shared signalling cascades or functional modules, thereby providing mechanistic insight into the underlying biological rationale of each prediction. This interpretability is crucial for translational applications, as it supports hypothesis generation for experimental validation and accelerates the prioritization of drug candidates with plausible biological mechanisms. Consequently, the proposed framework not only advances computational methodology but also strengthens the bridge between data-driven prediction and actionable biomedical discovery.

## Conclusions

In this study, we proposed a novel method, OMNI, a graph CNN model that integrates interaction information of different biomedical entities, including chemicals, diseases, genes, and pathways, to predict CGIs. It adopts a graph encoder-decoder architecture. The advantage of the OMNI model over other traditional graph neural network models is that, in the encoder part, it captures the heterogeneous network from both local and global views. Different decoders were evaluated, and an MLP that replaces the full dense weighted matrix with a low-rank factorized matrix. This decoder effectively decomposes the large layer into smaller ones, significantly reducing the number of parameters while preserving the information. The unique methodology incorporating both local and global view strategies, adopted in this study, outperformed the existing methods, as indicated by different evaluation results.

Notably, the introduction of more curated information results in a massive graph that is computationally challenging and exceeds the capability of available models. Thus, we developed a unique, suitable architecture for predicting multi-layer types. the OMNI adopts a DL framework to scale up the model’s performance and enable scalable computation in the training and prediction process. Further, the ability and the reliability of the models were evaluated in predicting high-quality results. Different case studies related to cancer and phytochemical-gene interactions were analyzed and demonstrated how OMNI can be applied to prioritise drug candidates by exploring the complex relationships related to the disease of interest. In the current work, we employed four types of entities, i.e. genes, chemicals, pathways, and diseases. In future work, we intend to expand the network by adding more types of entities, including symptoms and other biomedical data. The generated model with enriched data opens a new avenue in biomedical studies by predicting various biomedical relations.

## Data availability statement

All the data and source code underlying this article are available at https://github.com/NagaLab-MSN/OMNI. This also includes all the interactions predicted in two case studies and the supporting information.

## Acknowledgments

DBT is thanked for the financial support in the form of the Centre of Excellence in Advanced Computation and Data Sciences (Ref. No: BT/PR40188/BTIS/137/27/2021) and National Networking Project (NNP) (Ref. No: BT/PR40233/BTIS/137/74/2023). Manuscript communication number - CSIR-NEIST/PUB/2025/220.

## Biographical Note

**Gori Sankar Borah** holds a postgraduate degree in Computer Applications (MCA) and serves as a Scientific Administrative Assistant at CSIR-NEIST, with interests in machine learning, deep learning, and web development.

**Sukriti Tiwari** is a Junior Research Fellow at Mahindra University, India, working on brain–computer interfaces, bioinformatics, and machine-learning methods for biomedical signal processing and multimodal data analysis.

**Selvaraman Nagamani** is a Senior Scientist and Head of the Advanced Computation and Data Sciences Division (ACDSD), North East Institute of Science and Technology (NEIST), Jorhat, Assam, India. His current interests include Software Development, Machine Learning, Network Pharmacology.

